# Hsp90 dictates viral sequence space by balancing the evolutionary tradeoffs between protein stability, aggregation and translation rate

**DOI:** 10.1101/208462

**Authors:** Ron Geller, Sebastian Pechmann, Ashley Acevedo, Raul Andino, Judith Frydman

**Affiliations:** Department of Biology, Stanford University, Stanford, CA, USA; Department of Microbiology and Immunology, UCSF, San Francisco, CA, USA; Institute for Integrative Systems Biology (I2SysBio), Universitat de Valencia, Valencia, Spain; Department of Biochemistry, Université de Montréal, Montréal, QC, Canada

**Author notes:** Present Address: Laboratory of Virology and Infectious Diseases, Rockefeller University, Rockefeller University, New York, NY, USA. To whom correspondence should be addressed (R.A.), (J.F.). Equal contribution (Ron Geller, Sebastian Pechmann).

## Abstract

Acquisition of mutations is central to evolution but the detrimental effects of most mutations on protein folding and stability limit protein evolvability. Molecular chaperones, which suppress aggregation and facilitate polypeptide folding, are proposed to promote sequence diversification by buffering destabilizing mutations. However, whether and how chaperones directly control protein evolution remains poorly understood. Here, we examine the effect of reducing the activity of the key eukaryotic chaperone Hsp90 on poliovirus evolution. Contrary to predictions of a buffering model, inhibiting Hsp90 increases population sequence diversity and promotes accumulation of mutations reducing protein stability. Explaining this counterintuitive observation, we find that Hsp90 offsets the evolutionary tradeoff between protein stability and aggregation. Lower chaperone levels favor sequence variants of reduced hydrophobicity, thus decreasing protein aggregation propensity but at a cost to protein stability. Notably, reducing Hsp90 activity also promotes clusters of codon-deoptimized synonymous mutations at inter-domain boundaries, likely to promote local ribosomal slowdown to facilitate cotranslational domain folding. Our results reveal how a chaperone can shape the sequence landscape at both the protein and RNA levels to harmonize the competing constraints posed by protein stability, aggregation propensity and translation rate on successful protein biogenesis.

---

Acquisition of mutations is central to evolution but the detrimental effects of most mutations on protein folding and stability limit protein evolvability[1-4]. Molecular chaperones, which suppress aggregation and facilitate polypeptide folding[5, 6], are proposed to promote sequence diversification by buffering destabilizing mutations[7-10]. However, whether and how chaperones directly control protein evolution remains poorly understood. RNA viruses offer an attractive model to examine molecular mechanisms of evolution due to their relative simplicity and extreme evolutionary capacity. RNA viral polymerases generate mutations at almost the highest theoretically allowed rates[3, 11, 12]. The resulting genetic diversity of a viral population affords rapid adaptation to changing environments[13]. Recent findings link the high population diversity of RNA viruses to pathogenesis, likely due to the need to evolve rapidly in the infected organism[14]. Since viral proteins meet folding and stability challenges akin to those of host proteins, and utilize the host cell translation and folding machineries, viruses offer a unique opportunity to examine how chaperones shape protein evolution.

Poliovirus is a non-enveloped enterovirus that replicates in the cytosol of human cells. Infection leads to shutoff of host protein synthesis; thus the host translation and protein folding machineries are entirely dedicated to produce viral proteins[15]. Hsp90, an abundant ATP-dependent chaperone that facilitates the folding and maturation of many cellular metastable proteins[16-20], only interacts with a single poliovirus protein, the P1 capsid precursor[21]. Hsp90 activity is required for P1 folding and subsequent proteolytic maturation into capsid proteins VP0, VP1 and VP3[21]. Importantly, P1 folding is the only process in the poliovirus replication cycle that requires Hsp90. Pharmacological inhibition of Hsp90 activity with the highly selective Hsp90 inhibitor Geldanamycin (GA)[22] specifically impairs P1 folding and maturation but does not affect viral protein translation nor the folding or function of other viral proteins[21]. The high diversity and evolutionary capacity of poliovirus, together with the fact that it harbors only a single protein, the capsid precursor P1, that binds to and requires Hsp90 for folding, provide an ideal system to examine the role of Hsp90 on protein evolution (Fig. 1a).

**Figure 1.**
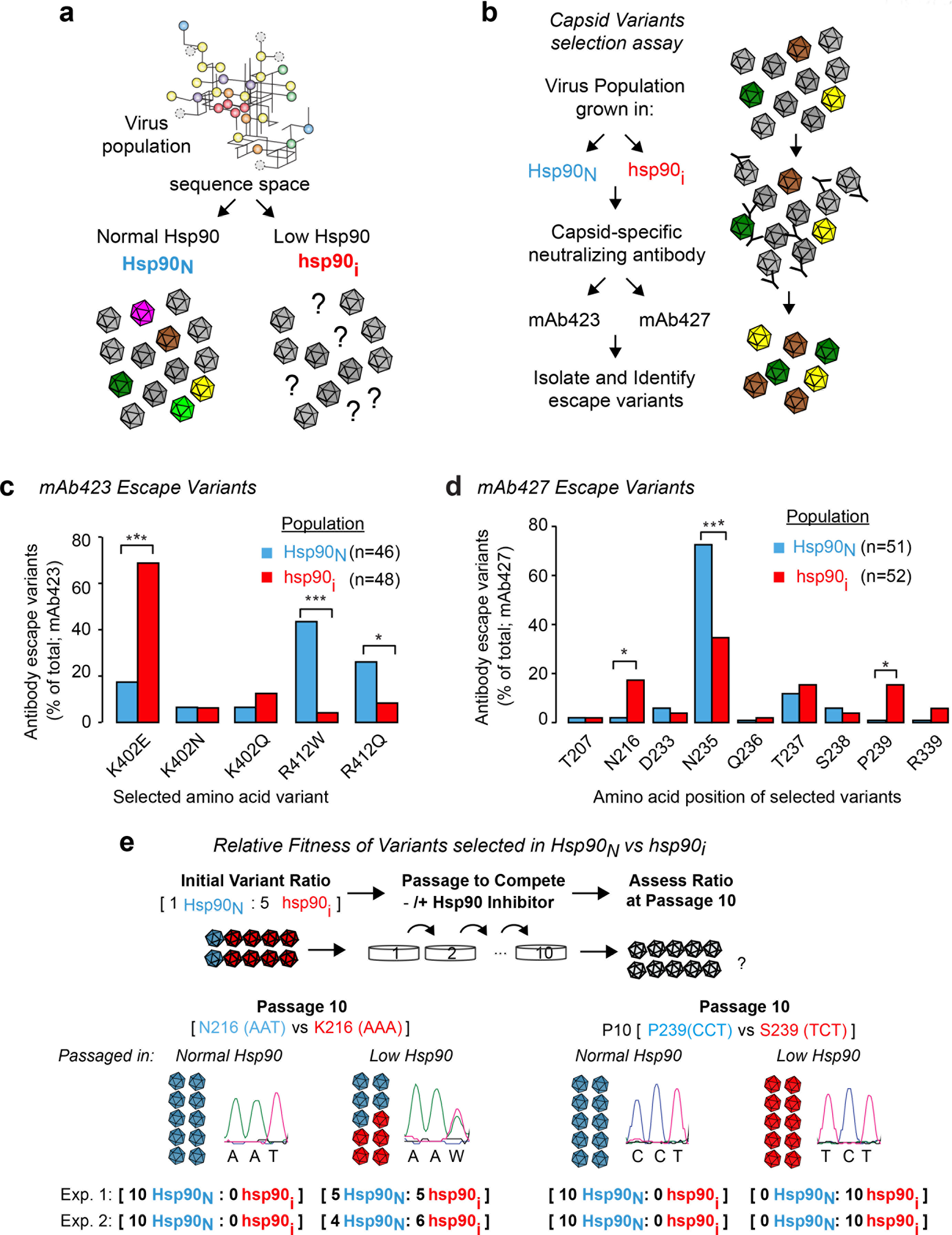
Hsp90 activity influences viral capsid fitness. (**a**) Does Hsp90 activity influence the sequence space of its client, poliovirus capsid protein P1? Schematic indicates a theoretical sequence space for P1 in viral populations, with each branch indicating two sequence variants linked by a point mutation. *Hsp90*_*N*_ and *hsp90*_*i*_ represent viral populations generated under normal Hsp90 activity, or under reduced Hsp90 activity by treatment with geldanamycin (GA). (**b**) Outline of capsid variants selection assay using neutralizing antibodies mAb423 and mAb427. (**c, d**) Abundance and distribution of mAb423 (c) or mAb427 (d) escape variants identified in *Hsp90*_*N*_ and *hsp90*_*i*_ populations. Each graph represents the sum of all mAb-resistant variants identified in two independent experiments (see Suppl. Fig. 1), with the total number of mutants (n) indicated. (e) Evaluation of the relative fitness of the *hsp90*_*i*_-enriched variants N216K and P239S identified in (d) under normal or Hsp90 inhibited conditions. An initial infection with a 1:5 ratio of *Hsp90*_*N*_ (N216 or P239) to *hsp90*_*i*_ variant (K216 or S239) virus was used, and viruses serially passaged under normal or Hsp90 inhibited conditions for 10 passages. The results of the competition assays were assessed by Sanger sequencing of the population, and is shown for passage 10 (P10). * p < 0.05, ** p < 0.01, *** p < 0.005 by two-tail Fisher’s exact test after multiple test correction using the FDR method.

During infection, viral capsids are subject to intense selective pressure by the host immune system[23, 24]. Viral protein diversity is key to virus survival as it creates a reservoir of capsid variants enabling escape from neutralization by circulating antibodies. To assess whether Hsp90 influences viral capsid evolution, we examined the spectrum of neutralizing antibody escape variants for viral populations that were generated under normal conditions (*Hsp90*_*N*_) or following Hsp90 inhibition by treatment of infected cells with GA (*hsp90*_*i*_) (Fig. 1b). Two neutralizing monoclonal antibodies (mAb423 and mAb427) targeting different epitopes in the poliovirus capsid[25] were employed, followed by sequencing across their respective epitope. Escape variants conferring resistance to neutralization were identified in both conditions, in two independent experiments (Fig. 1b). For mAb423, resistance in the *Hsp90*_*N*_ population mapped to several different substitutions in two P1 sites: K402 and R412 (Fig. 1c, Suppl. Fig. 1a, Suppl. Table 1). In contrast, mAb423 resistance in the *hsp90*_*i*_ population was dominated by a single substitution, K402E (p < 0.0001 by Fisher’s test; Fig. 1c, Suppl. Fig. 1a, Suppl. Table 1). The R412W mutation, often encountered in the *Hsp90*_*N*_ population, was significantly disfavored in the *hsp90*_*i*_ population (p < 0.0001 by Fisher’s test; Fig. 1c, Suppl. Fig. 1a, Suppl. Table 1).

For the other antibody, mAb427, escape neutralization variants observed in both *Hsp90*_*N*_ and *hsp90*_*i*_ populations were distributed across multiple sites in the epitope. Several distinct variants at position 235 (N235) were significantly enriched in the *Hsp90*_*N*_ population (p = 0.0014 by Fisher’s test; Fig. 1d, Suppl. Fig. 1b, Table 2). In contrast, mutations at positions N216 and P239 were observed at high frequency in the *hsp90*_*i*_ population (>15%) but were nearly absent from the *Hsp90*_*N*_ population (<2%; p < 0.05 by Fisher’s test for both N216 and P239; Fig.1d, Suppl. Fig. 1b, Suppl. Table 2).

These results suggest that Hsp90 dictates the sequence space of P1 by selecting both for and against specific sequence variants. The near absence from the *Hsp90*_*N*_ population of mAb427 escape variants at sites N216 and P239 suggests that normal Hsp90 levels reduce the fitness of these sequence variants. This finding deviates from the buffering model of chaperone action, whereby Hsp90 is proposed to generally buffer detrimental mutations in client proteins[10, 26-31]. To directly test the notion that Hsp90 negatively selects certain sequence variants, we assessed the relative fitness of viruses harboring either the *hsp90*_*i*_ selected variants at P1 sites 216 and 239, or the *Hsp90*_*N*_ variants observed under Hsp90-normal conditions. Viruses containing the *Hsp90*_*N*_ sequences N216 and P239 were competed with viruses containing the *hsp90*_*i*_ variants K216 or S239, under conditions of either normal or low Hsp90 activity (Fig. 1e). A mix with an initial ratio of 1:5 *Hsp90*_*N*_ to *hsp90*_*i*_ variant was serially passaged ten times with or without Hsp90 inhibitor in two independent experiments. Sequence analysis of passage 10 populations (P10, Fig. 1e and Suppl. Fig. 2) showed that under normal Hsp90 conditions, the *Hsp90*_*N*_ variants N216 and P239 were strongly selected for in both independent experiments, completely dominating the population after 10 passages despite the initial excess of *hsp90*_*i*_ virus (Fig. 1e, Normal Hsp90). In contrast, when passaged in the presence of Hsp90 inhibitor (Fig. 1e, Low Hsp90), *hsp90*_*i*_ variant K216 was present in ~50% of the population (Fig. 1e and Suppl. Fig. 2a) and variant S239 completely dominated the population (Fig. 1e and Suppl. Fig. 2b). These findings demonstrate that Hsp90 levels dictate the fitness of particular sequence variants in a client protein. Given the growing interest in Hsp90 inhibitors as broad-spectrum antivirals[21, 32, 33], the effect of Hsp90 on the fitness of escape mutants from immune selection may have important therapeutic implications.

To gain a better understanding of how Hsp90 shapes virus evolution, we next examined the diversity of poliovirus populations at unprecedented resolution using recent technological advancements in ultra-deep sequencing that provide a detailed description of the viral population mutation composition[34]. We considered three possible scenarios by which Hsp90 influences the functional sequence space of a protein (Fig. 2a). The first possibility, that Hsp90 plays no role shaping sequence space (*No Effect*, Fig. 2a, *i*), is negated by the experiments in Fig. 1. The second model stems from the role of Hsp90 and other chaperones in buffering destabilizing mutations[10, 26-31] (*Buffers Mutations;* Fig. 2a, *ii*). This *buffering* model predicts that Hsp90 inhibition will subject less stable mutations to purifying selection due to constraints imposed by protein biophysics[35, 36]. This should favor a smaller sub-population of stable variants, resulting in reduced population diversity. Finally, a third possible model is that Hsp90 shapes sequence space in a hitherto unanticipated manner (*Dictates Landscape*, Fig. 2a *iii*).

**Figure 2.**
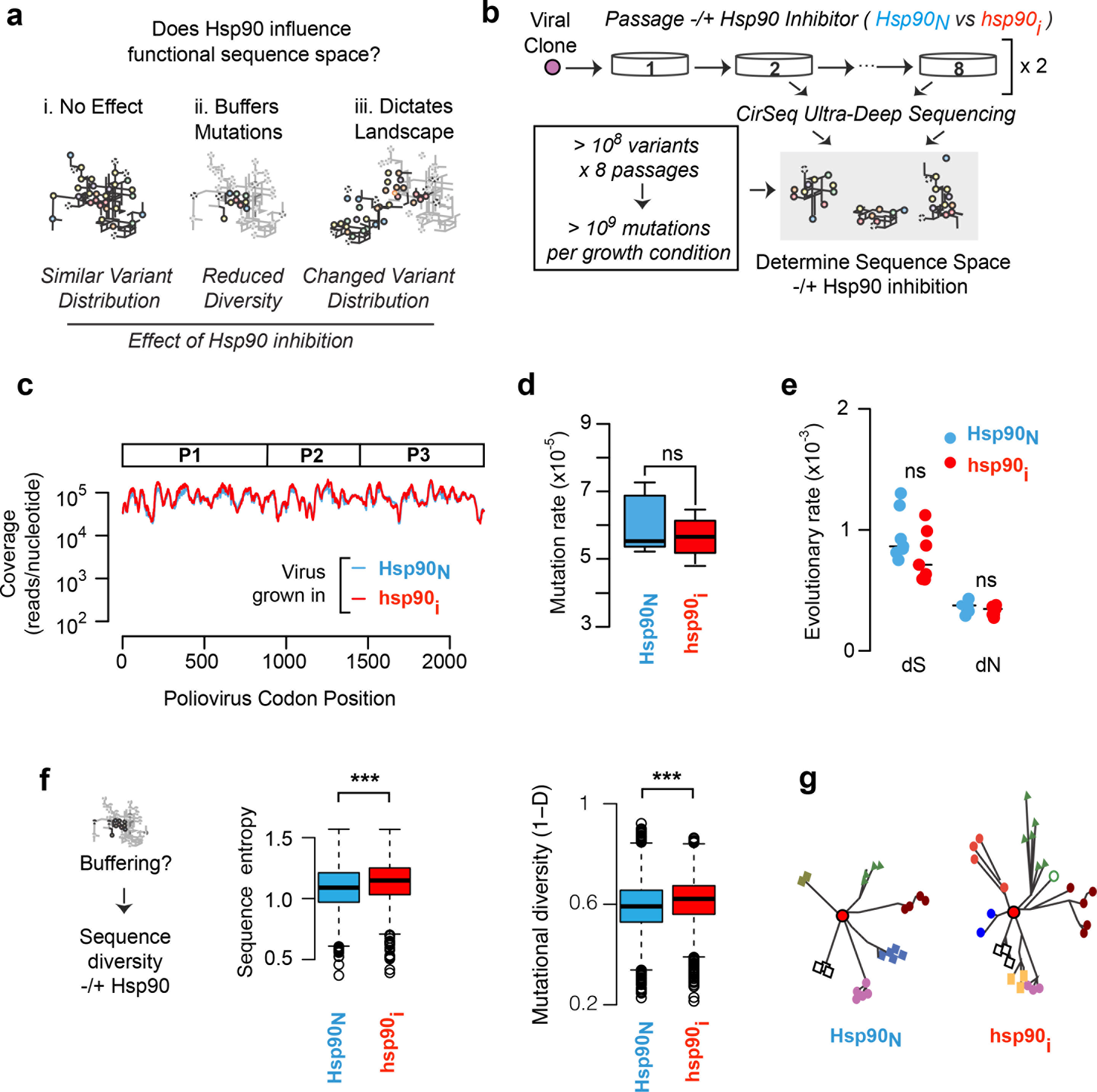
CirSeq deep sequencing of poliovirus populations to globally assess the effects of Hsp90 on protein sequence space. (**a**) Hypothetical models for how Hsp90 influences the functional sequence space of poliovirus: *i*. No effect, *ii*. Buffers mutations, or *iii*. Dictates landscape. Schemes illustrate the respective predicted outcome of Hsp90 inhibition on sequence space for each model. (**b**) Experimental strategy to examine the effects of Hsp90 inhibition on poliovirus sequence space. Poliovirus was propagated over 8 passages under normal or low Hsp90 conditions via GA treatment. Population samples containing >10^8^ variants each were collected from passages 2-8 and subjected to CirSeq deep sequencing and analysis. (**c**) Average sequence coverage obtained with CirSeq across the poliovirus genome for all populations *Hsp90*_*N*_ (blue trace) and *hsp90*_*i*_ (red traces). (**d**) Mutation rates for the full virus genome in *Hsp90*_*N*_ and *hsp90*_*i*_ populations. (**e**) evolutionary rates for P1 in *Hsp90*_*N*_ and *hsp90*_*i*_ populations (full genome in Suppl. Fig. 4c) (**f**) Does Hsp90 buffering lead to reduced P1 sequence diversity in *hsp90*_*i*_ populations? Sequence entropy (Left panel) and Mutational Diversity (Right panel) for P1 coding region for is higher for *hsp90*_*i*_ than for *Hsp90*_*N*_ populations analyzed by CirSeq. (**g**) Representation of distinct sequence diversity for P1 from *Hsp90*_*N*_ and *hsp90*_*i*_ virus populations. The original master sequence (red, middle circle) gives rise to variants (different colors and symbols) through mutation. Significance: ns, not significant (p > 0.05); *** p < 0.05 by Wilcoxon-test.

Starting with a single viral clone, we sampled virus populations from eight different passages under conditions of normal Hsp90 activity (*Hsp90*_*N*_) or low Hsp90 activity (*hsp90*_*i*_) by treatment with the Hsp90 inhibitor GA[22] (Fig. 2b) in two independent replicates (Fig. 2b). We then employed high fidelity CirSeq ultradeep sequencing[34] to define the sequence composition of these poliovirus populations (Fig. 2b). Extensive sequence coverage was obtained for both replicas, with most sites in the coding sequence covered over 100,000 times (Fig. 2c and Suppl. Fig. 4a). The consensus sequence was unchanged in all samples, as expected from previous findings that poliovirus cannot gain resistance to Hsp90 inhibitors[21]. Furthermore, no global differences in the overall mutation rates of the *Hsp90*_*N*_ and *hsp90*_*i*_ virus populations were observed (Fig. 2d, p = 0.71 by Mann-Whitney-Wilcoxon (MWW) test, and Suppl. Fig. 4b). Similarly, no significant differences in overall evolutionary rates were observed for both synonymous (*dS*) and non-synonymous mutations (*dN*) as a function of Hsp90 activity level, either for Hsp90 client protein P1, or across the entire genome (Fig. 2e, dS: p = 0.21 and dN: p = 0.25 by MWW test; and Suppl. Fig. 4c). Thus, global evolutionary rates were not affected by Hsp90 activity, consistent with previous findings that Hsp90 inhibition does not affect the viral polymerase[21]. Interestingly however, protein mutational diversity in P1 was significant higher in the *hsp90*_*i*_ condition, in contrast to the expectation from the *buffering* model (Fig. 2f). Two alternative metrics, sequence entropy (Fig. 2f, p < 8×10^-26^ by MWW test; and Suppl. Fig. 4d) and mutational diversity (1-D) with D as Simpson’s index of diversity (Fig. 2f, p < 8.7 ×10^-21^ by MWW test; and Suppl. Fig. 4d) both indicated that reducing Hsp90 activity enhances population diversity (Fig. 2g). Together with the data from Fig. 1, these analyses suggest that Hsp90 dictates the sequence landscape within the population (Fig. 2a, model *iii*).

Molecular chaperones assist protein homeostasis by promoting folding upon translation, maintaining stability and also preventing protein aggregation[5, 37, 38] (Fig. 3a). These processes are particularly relevant to viral capsids, which must be produced in high levels and assembled from numerous subunits, while avoiding aggregation[39-42]. Furthermore, once assembled, capsids must be sufficiently stable to protect the viral genome in harsh environments. To further understand the role of Hsp90 in shaping protein evolution, we compared the viral populations generated under normal (*Hsp90*_*N*_) or Hsp90-inhibited (*hsp90*_*i*_) conditions to identify variants that differed significantly in their effect on protein stability or aggregation propensity (Fig. 3b). First, we calculated the effects of all possible single point mutations on protein stability in the three viral proteins for which crystal structures are available, the capsid P1, the protease 3C and the RNA polymerase 3D (FoldX algorithm[43] see Methods and Suppl. Fig. 5a). We then compared *Hsp90*_*N*_ and *hsp90*_*i*_ populations to identify sites that significantly differ in their destabilizing variant frequencies (*Site*_*destab*_). Next, we calculated the effect of mutations on sequence aggregation propensity (TANGO algorithm[44] see Methods and Suppl. Fig. 5b) in these same viral proteins to uncover sites that differ in their ability to tolerate variants of increased aggregation propensity (*Sites*_*agg*_) between the *Hsp90*_*N*_ and *hsp90*_*i*_ population. Surprisingly, there were significantly more *Site*_*destab*_ in Hsp90 client P1 under low Hsp90 activity (Fig. 3c and Suppl. Fig. 5d; p = 0.0009 by Cochran–Mantel–Haenszel (CMH) test). Conversely, the number of *Sites*_*agg*_ in P1 was significantly reduced under low Hsp90 activity (Fig. 3d, p = 0.00036 by CMH test). The incidence of *Sites*_*agg*_ and *Site*_*destab*_ in the non-Hsp90 dependent viral proteins 3C and 3D was relatively unaffected by Hsp90 inhibition (Fig. 3c, 3d: p = 0.21, 3D: p = 1 by CMH test). These analyses indicate that Hsp90 preferentially affects the functional sequence space of its client P1. Paradoxically, unlike what we expected from the ‘*buffering*’ model, reducing Hsp90 activity increases the incidence of variants harboring destabilizing mutations and favors variants that reduce aggregation propensity, in support of a ‘*dictating landscape*’ model.

**Figure 3.**
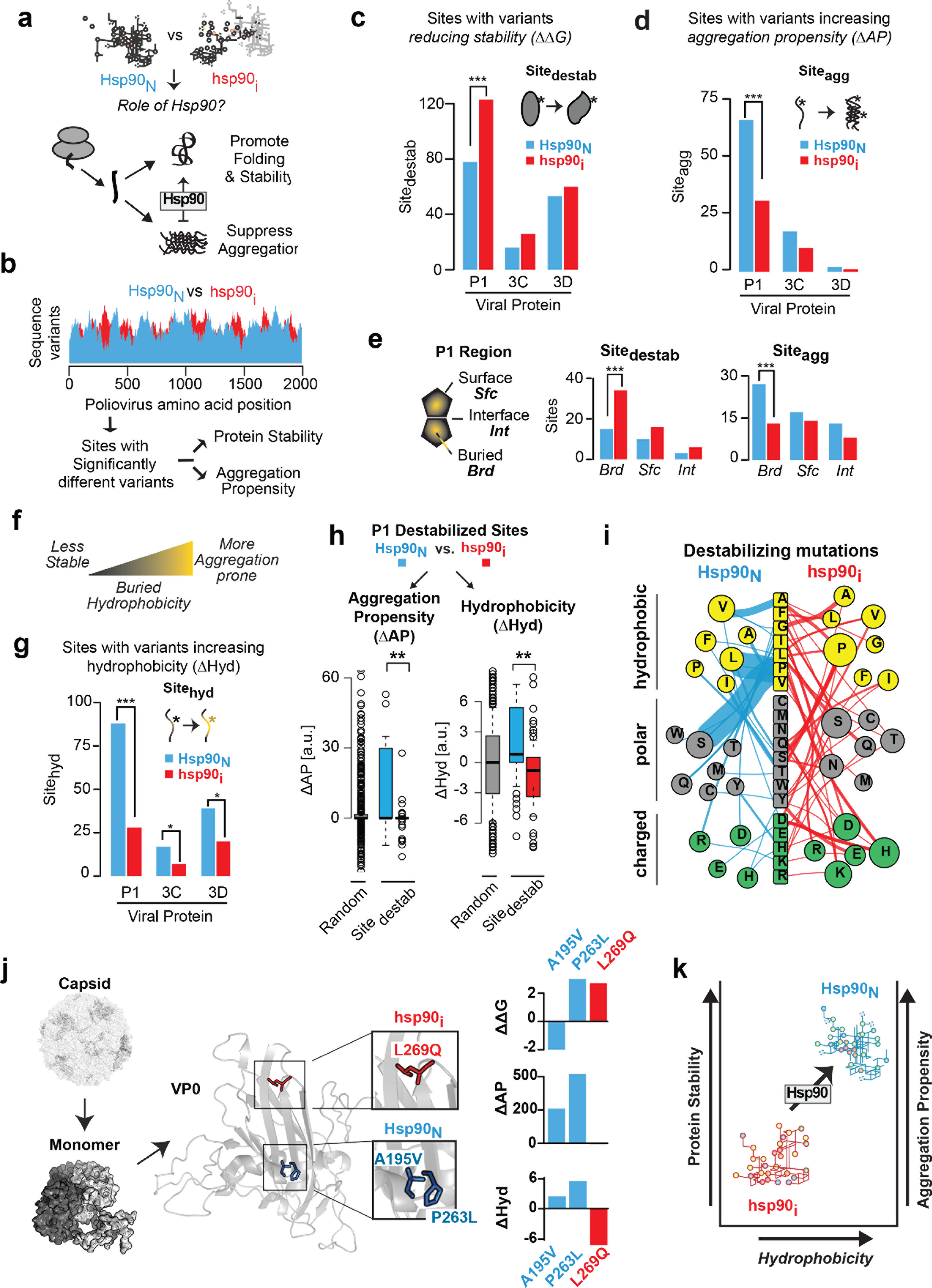
Hsp90 influences the capsid sequence space. (**a**) Role of Hsp90 in pathways of protein folding and aggregation. (**b**) Analysis pipeline to identify codon sites that differ significantly in their ability to support destabilizing variants (Site_destab_) or aggregation prone (Site_agg_) variants between *Hsp90*_*N*_ and *hsp90*_*i*_ populations. Significant sites were identified as described in Methods. (**c, sd**) Number of Site_destab_ (**c**) and Site_agg_ (**d**) observed in viral proteins: P1, capsid protein; 3C, protease; and 3D, polymerase. (**e**) Distribution of Site_destab_ and Site_agg_ at buried (Brd), interface (Int), or surface (Sfc) positions across folded P1 (based on crystal structure 2PLV). (**f**) Effect of buried hydrophobicity on protein stability and aggregation propensity (**g**) Sites with increased hydrophobic variants (Site_hydro_) in *Hsp90*_*N*_ and *hsp90*_*i*_ populations. (**h**) Properties of P1 Destabilized sites (Site_destab_) from *Hsp90*_*N*_ and *hsp90*_*i*_ aggregation propensity (left panel) and hydrophobicity (right panel). (**i**) Distinct nature of destabilizing variants (Site_destab_) in P1 of *Hsp90*_*N*_ and *hsp90*_*i*_ populations. Original amino acid is indicated in the middle. Colored indicates side-chain properties: Yellow: hydrophobic; Gray: polar; Green: charged. Mutations to specific residues in *Hsp90*_*N*_ (left) and *hsp90*_*i*_ (right) are indicated, where circle size indicates mutation occurrence. The width of lines connecting each mutation represents the number of occurrences for each variant. (**j**) Exemplar P1 variants enriched in *Hsp90*_*N*_ and *hsp90*_*i*_ populations. Location in P1 folded structure and effects on stability, aggregation propensity, and hydrophobicity are indicated. Mutations A195V and P263L (blue) observed in *Hsp90*_*N*_, and L269Q (red) in *hsp90*_*i*_. Structure from 2PLV, with VP2 monomer, and assembled capsid indicated. (**k**) Schematic effect of Hsp90 on sequence space. Hsp90 allows variants to explore a region in sequence space of increased hydrophobicity and aggregation propensity, but also of increased stability. Lower Hsp90 constrains sequence space variants to a region of reduced stability, hydrophobicity, and aggregation propensity. Significance levels are indicated as * p < 0.05; ** p < 0.01; *** p < 0.005.

We next examined the distribution of *Sites*_*agg*_ and *Site*_*destab*_ with respect to the folded P1 structure, namely whether they map to buried (*Brd*), surface exposed (*Sfc*) or interface (*Int*) positions (Fig. 3e) For both *Sites*_*agg*_ and *Site*_*destab*_, major differences between *Hsp90*_*N*_ and *hsp90*_*i*_ mapped to buried (*Brd*) positions within the structure of the viral capsid (Fig. 3e and Suppl. Fig. 5d). Under low Hsp90 conditions, buried (*Brd*) destabilizing variants were increased (*Site*_*destab*_; p = 0.006 by Fisher’s test) and aggregation prone sites decreased (*Sites*_*agg*_; p < 0.004 by Fisher’s test) compared to surface exposed (Fig. 3e; *Sfc*) or interface regions (*Int*, Fig. 3e and Suppl. Fig. 5e). Since buried regions are primarily exposed in non-native conformations, such as those generated during cotranslational folding, these findings indicate that Hsp90 activity may be critical to prevent aggregation during initial protein folding, rather than during assembly, for which interface residues are likely of greater relevance.

Disfavoring aggregation-prone variants should be beneficial for the virus, while favoring destabilizing mutations should be detrimental. How can the contradictory effects of low chaperone activity on these two key protein-folding properties be reconciled? We reasoned that hydrophobicity, a driving force in protein folding that influences both protein stability and aggregation[45, 46] may explain these effects (Fig. 3f). Enhancing buried hydrophobicity can increase the stability of the folded core, but can also increase aggregation propensity. We thus examined if *Hsp90*_*N*_ and *hsp90*_*i*_ populations differ in their ability to accommodate sites with enhanced hydrophobicity (*Sites*_*hydro*_; Fig. 3g). Strikingly, inhibiting Hsp90 significantly reduced the number of *Sites*_*hydro*_ in P1 and to a lesser extent also in non-Hsp90 clients 3C and 3D (Fig. 3g). The effect on P1 was dramatic for both replicas (Fig. 3g, p = 5×10^-9^ by CMH test, and Suppl. Fig. 5f) and observed both in buried, surface-exposed and interface sites (Suppl. Fig. 5c, *Brd*: p = 0.0037, *Int*: p = 0.0033, *Sfc*: p = 0.018 by Fisher’s test). These results indicate that Hsp90 enables sequence variants of increased hydrophobicity, which can promote protein stability but also increase aggregation propensity.

To better understand the surprising finding that reduced chaperone function results in an increase in destabilizing variants, we next examined the properties of individual *Site*_*destab*_ variants in *Hsp90*_*N*_ and *hsp90*_*i*_ viral populations. *Site*_*destab*_ variants in P1 were examined with respect to their ability to change aggregation-propensity (ΔAP) and hydrophobicity (ΔHyd) (Fig. 3h,i). Destabilizing variants observed under normal Hsp90 conditions (*Hsp90*_*N*_) were significantly more aggregation-prone than destabilizing variants present at reduced chaperone activity (*hsp90*_*i*_; Fig. 3h, p = 0.002 by MWW test, and Suppl. Fig. 5g). Furthermore, *Site*_*destab*_ variants in *hsp90*_*i*_ were significantly less hydrophobic than in *Hsp90*_*N*_ or those expected by chance (i.e. randomly selected mutations; Fig. 3h, p< 10^-7^ by MWW test, and Suppl. Fig. 5h). This indicates that *Hsp90*_*N*_ and *hsp90*_*i*_ accommodate different kinds of destabilizing mutations. Thus, destabilizing mutations in *Hsp90*_*N*_ have significantly increased sequence hydrophobicity and aggregation propensity compared to that expected by chance (Fig. 3h, left panel, p < 10<^-5^ by MWW test and Suppl. Fig. 5g). In contrast, destabilizing mutations in *hsp90*_*i*_ have significantly decreased sequence hydrophobicity and aggregation propensity (Fig. 3h, right panel, p = 0.0007 by MWW test, and Suppl. Fig. 5g). More specifically, many destabilizing mutations in the *Hsp90*_*N*_ population resulted from the loss of proline residues (Fig. 3i and Suppl. Fig. 5h). In contrast, destabilizing variants in the *hsp90*_*i*_ population were more diverse and reduced hydrophobicity by mutation to polar or charged residues (Fig. 3i and Suppl. Fig. 5h). We conclude that Hsp90 enables accumulation of destabilizing variants with increased aggregation propensity and hydrophobicity, while reducing Hsp90 activity promotes destabilizing variants that reduce hydrophobicity and aggregation propensity.

The interplay between Hsp90 action and individual mutations affecting protein stability, aggregation propensity and hydrophobicity is illustrated by a set of variants found at the interface between core beta strands in VP0, the N-terminal folding unit of P1 (Fig. 3j). For instance, a stabilizing mutation found in *Hsp90*_*N*_, A195V, should be beneficial to protein folding and stability; however, this mutation also increases P1 aggregation propensity and hydrophobicity, and is thus strongly selected against in the *hsp90*_*i*_ population (p < 1×10^-4^ by CMH test). Another *Hsp90*_*N*_ variant, P263L, also increases aggregation propensity and hydrophobicity but, in this case, reduces protein stability. This very detrimental mutation is strongly depleted under *hsp90*_*i*_ conditions (p < 1×10^-4^ by CMH test). In contrast, L269Q, which reduces sequence hydrophobicity and does not alter aggregation propensity, is favored in *hsp90*_*i*_, despite being a strongly destabilizing mutation (p < 10^-10^ by CMH test). These analyses further illustrate that, for poliovirus P1 capsid protein, the dominant selective force under low Hsp90 activity is the reduction of hydrophobicity and aggregation propensity, even at a cost to protein stability.

Our analysis exposes the role of Hsp90 in balancing an intrinsic conflict between stability and aggregation, shaping protein evolution (Fig. 3k). While increased hydrophobicity benefits the stability of the folded state, it also increases the aggregation propensity of folding intermediates, such as those generated during protein synthesis. Thus, chaperone-dependence, which reduces aggregation *in vivo*, may allow exploration of regions within sequence space leading to more stable, and thus more versatile, proteins (Fig. 3k). This is consistent with the high aggregation propensity of obligate chaperone substrates in the cell[47, 48]. Importantly, by favoring more stabilizing but also more aggregation-prone sequence variants, chaperones appear to restrict the sequence space and render proteins vulnerable to aggregation under conditions of chaperone impairment, such as cellular stress or aging.

Our analysis points to the importance of Hsp90 in preventing aggregation during initial protein folding (Fig. 3e). The vectorial nature of translation renders proteins particularly susceptible to aggregation[49], since emerging domains that cannot yet fold will expose hydrophobic elements to the crowded cellular milieu[50]’[51]. Importantly, chaperones play a key role protecting nascent chains and facilitating their folding[52]. Since the speed of translation is proposed to be directly attuned to optimize the cotranslational folding and processing of nascent chains[53-57], we hypothesized that virus adaptation to low Hsp90 levels may involve optimization of the translation elongation rate. Some tRNAs are much more abundant than others and codon choice can have important effects on translation kinetics[58-60] (Fig. 4a). Accordingly, viruses have been found under direct selection for their synonymous codon usage[61] whose deregulation often brings direct fitness consequences[62-64]. We thus examined whether reducing Hsp90 levels affected the synonymous choice of codons in P1 using the established tRNA adaptation index (tAI)[65] for both the human genome and for HeLa cells used for virus passaging. Of note, the tAI has been shown to correlate with *in vivo* translation speed[59]. Overall, both *Hsp90*_*N*_ and *hsp90*_*i*_ populations demonstrate a clear trend to generally optimize overall codon usage to match that of the host cell (Fig. 4b; *Hsp90*_*N*_: p < 10^-6^ by MWW test; *hsp90*_*i*_: p < 10^-6^; Suppl. Fig. 6a,b). This indicates that poliovirus adapts to the host cell to enhance overall protein production, establishing a direct link between viral infection and adaptation to the host cell tRNA pool.

**Figure 4.**
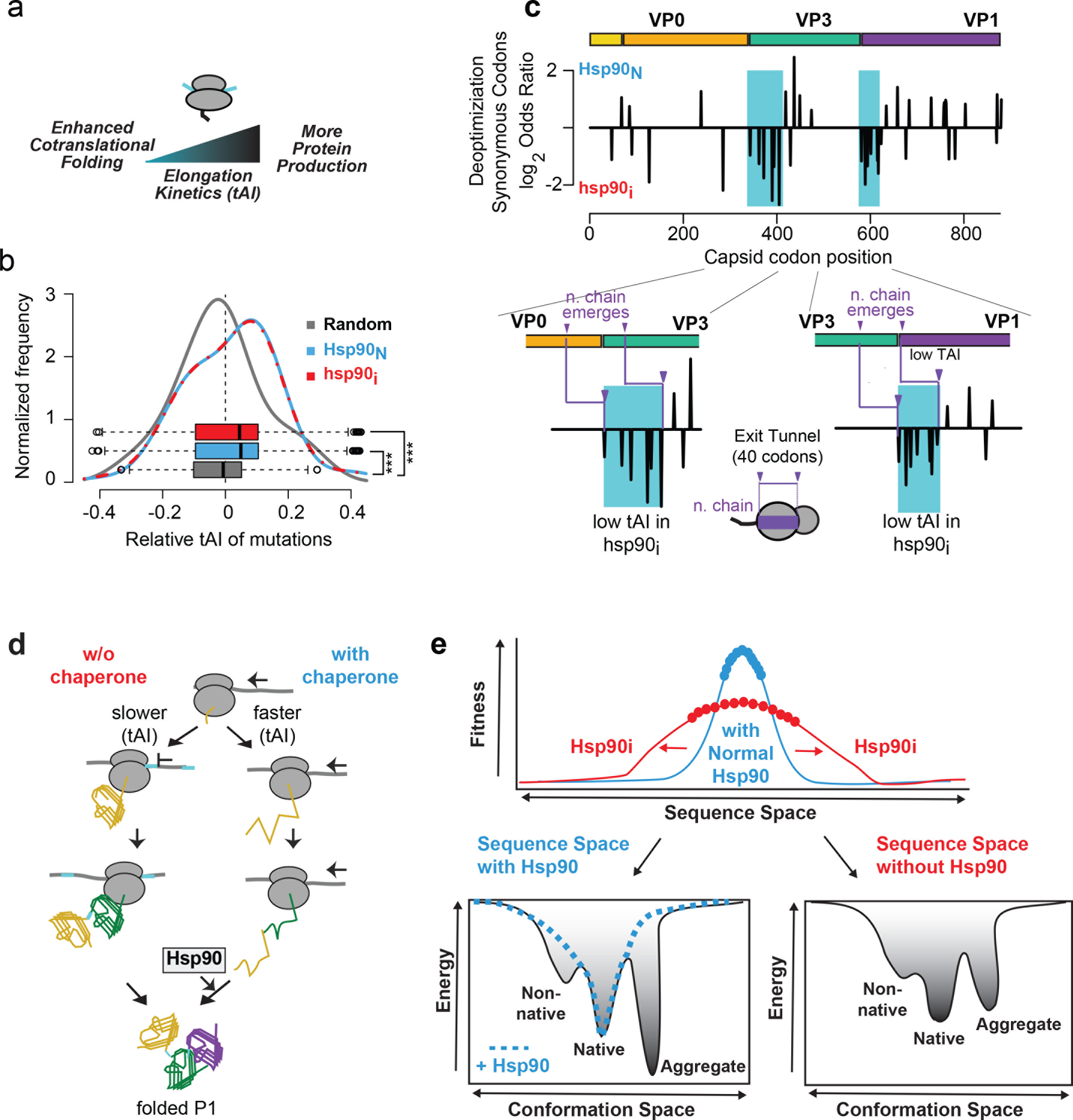
Codon deoptimization at domain boundaries upon adaptation to Hsp90 inhibition. (**a**) Interplay between the rate of translation, cotranslational folding and protein production. (**b**) Both *Hsp90*_*N*_ and *hsp90*_*i*_ poliovirus populations accumulate synonymous variants resulting in global codon optimization. The change in codon optimality was calculated for all synonymous mutations observed in *hsp90*_*i*_ and *Hsp90*_*N*_ populations as well as for a random sample of mutations (random) based on the hela cell tRNA adaptation index (tAI). (**c**) Distribution of sites harboring significantly deoptimized synonymous codons across P1 in *hsp90*_*i*_ and *Hsp90*_*N*_ populations. The domain structure of P1 is indicated. Insets show the two clusters of deoptimizated codons in *hsp90*_*i*_ (indicated with blue shading), relative to domain boundaries of VP0/VP3 (left) and VP3/VP1 (right). The approximate size of the ribosomal exit tunnel size and the point at which the nascent chain (n. chain) emerged from the ribosome are indicated in purple (**d**) Model of the potential impact of chaperone levels on codon deoptimization at domain boundaries. Under conditions of Hsp90 inhibition, the virus locally reduces translation speeds to promote cotranslational folding. While enhancing correct cotranslational P1 folding, this strategy may slow protein production. (**e**) Role of Hsp90 in shaping protein evolution by modulating the energy landscape of protein folding and balancing trade-offs between protein stability, aggregation propensity and folding. Hsp90 allows exploration of variants that increase aggregation propensity and hydrophobicity, which may also lead to increased protein stability. Under reduced Hsp90 conditions, the viral sequence space expands to explore variants that reduce aggregation propensity at the expense of stability. See text for details.

However, under conditions of Hsp90 inhibition, we also observed clusters of sites harboring synonymous mutations that deoptimized codons at specific locations along the P1 coding sequence (Fig. 4c and Suppl. Fig. 6c,d). Specifically, there were two major clusters of deoptimized codons in the *hsp90*_*i*_ population (Fig. 4c, highlighted in blue). Each cluster mapped to the N-terminal coding region of VP3 and VP1, respectively. The P1 polyprotein encompasses three domains that, upon completion of folding, are processed proteolytically to generate processed capsid subunits VP0, VP3 and VP1[40]. Interestingly, accounting for the length of the ribosomal exit tunnel, the deoptimized codon clusters were positioned to promote a local slow-down of translation elongation upon emergence of the complete VP0 and VP3 folding domains from the ribosomal exit site (Fig. 4c, bottom panels). Importantly, pulse chase experiments show that Hsp90 inhibition does not affect global translation speeds in poliovirus infected cells[21], supporting the idea that synonymous changes affect the translation rate locally. We propose that synonymous mutations are selected at lower Hsp90 activity to locally reduce translation speed at inter-domain boundaries to favor co-translational folding. Consistent with this notion, the only member of the picornavirus family whose P1 protein does not require Hsp90 for folding, Hepatitis A virus, relies on highly deoptimized codon-usage for the capsid coding region for fitness[66]. These findings suggest that achieving overall higher translation rates also requires a tradeoff with efficient cotranslational folding, which can be balanced by chaperone action or in its absence, by selective codon deoptimization at specific sites (Fig. 4d).

While clearly essential for protein folding[16], the role of Hsp90 in evolution has remained unclear[67]. Our study demonstrates that the chaperone Hsp90 helps shape the evolutionary path of a viral protein at the polypeptide and RNA levels. Hsp90 alters the protein sequence space by modulating tradeoffs between protein stability, aggregation and hydrophobicity at the polypeptide level, and the tradeoffs between translation speed and cotranslational folding at the RNA level. Hsp90 allows exploration of variants that increase aggregation propensity and hydrophobicity, which may also lead to increased protein stability (Fig. 4e). In low Hsp90, the viral sequence space expands to explore variants that reduce aggregation propensity at the expense of stability.

Our experiments provide insights into how chaperones may have shaped protein evolution by modulating the energy landscape of protein folding. Upon synthesis, many globular proteins populate non-native states as well as aggregation prone-intermediates; of note, aggregates are often more stable than the native folded state[68] (Fig. 4e). Chaperones guide the folding polypeptide through this landscape to prevent aggregation and favor the energetically stable folded conformation. Without chaperones, alternative solutions within sequence space must be explored that alleviate the pressure to reduce intrinsic aggregation propensity; this may preclude the emergence of more hydrophobic variants that stabilize the native state. These considerations may be particularly relevant in the crowded environment of the cell, as well as for topologically complex proteins such as viral capsids, which must assemble elaborate oligomeric structures[69]. However, the intrinsic conflict between protein stability and aggregation propensity is a property of all proteins[68] and we speculate that our results have general implications for further understanding how chaperones shaped the evolution of proteins.

Our experiments also suggest that chaperone activity influences the RNA sequence space. Low Hsp90 promotes deoptimizing codons following translation of domains. Such modulation of local translation kinetics may enhance cotranslational folding by allowing more time for early folding intermediates to adopt correct conformations, avoid misfolding and aggregation, or by allowing more time for recruiting and interacting with chaperones. These findings highlight the close relationship between protein and RNA sequence space, and show that the evolutionary trajectory of both may be shaped by chaperones.

Not all proteins require chaperone assistance for folding[70]. Our results suggest that one benefit of acquiring chaperone-dependence is to resolve evolutionary trade-offs that allow proteins to be translated faster and to occupy specific regions of sequence space. Chaperone dependence thus likely supported acquisition of complex folds that expand protein function, including meta-stable structures[19, 20], multimeric assemblies[5, 71], and multi-domain architectures[5, 71]. On the other hand, chaperone substrates are constrained by chaperone-specific preferences and become sensitive to the levels and activity of that chaperone in different cells and environmental conditions. Chaperone activity can vary with stress, cell type and even age[37, 72], and these changes may yield variance in the folding and activity of a chaperone-dependent protein. In the particular case of viral proteins, where survival and adaptation is directly dependent on population diversity [14], chaperones likely play a fundamental role in virus protein evolution, adaptation to specific environments, like tissues and stress conditions, and ultimately pathogenesis.

## Methods

### Cells, viruses and reagents

All experiments were conducted in HeLa S3 cells cultured in DMEM/F12 (Invitrogen) containing 10% FCS, penicillin and streptomycin and at 37°C. The Mahoney type 1 (WT) and Sabin type 1 strains of poliovirus were generated, and infections carried out, as previously described[21]. Geldanamycin (GA; LC labs) was diluted in DMSO. Monoclonal poliovirus antibodies were generously provided by Dr. Morag Ferguson at NIBSC, Blanche Lane, South Mimms, Potters Bar, Hertfordshire.

### Antibody neutralizations, and competition assays

For selection of antibody escape mutants, ~10^5^ (mAb427) or 10^6^-10^7^ (mAb 423) PFU (Plaque Forming Units) from virus populations generated in the absence or presence of GA (as described above) were mixed with 1μL of antibody for 1 hour at 25 °C and then used to infect confluent cells in 6 cm plates for 30 minutes. Cells were then washed and overlaid with DMEM/F12 containing 1% agar and 1:2000 dilution of antibody. Twenty-four to 30 hours later, plaques were picked and amplified by inoculating onto fresh cells in 12 well plates. Upon CPE, viral RNA was extracted by Trizol or ZR Viral RNA kit (Zymo Research), RT-PCR performed with Thermoscript reverse transcriptase (Invitrogen) using random primers, and PCR performed using primers F676 (5’-GCTCCATTGAGTGTGTTTACTCTA-3’) and R3463 (5’-ATCATCCTGAGTGGCCAAGT-3’) and PfuUltra II Fusion HS (Stratagene). For competition experiments, 10^5^ PFU of mutants were mixed with 2×10^4^ pfu of WT poliovirus and passaged as above. P-values were calculated using a two-tailed fisher’s test and multiple testing correction using the FDR method[74].

### CirSeq analysis

CirSeq was performed according to published protocols[73], with the exception of the presence of 1 μM Hsp90 inhibitor Geldanamycin during the evolution of the *hsp90*_*i*_ condition. The virus mutation rate was computed by analyzing mutations to stop codons as described previously[75]. The evolutionary rates *dN* and *dS* were computed with established count model of codon sequence evolution by Nei & Gojobori[76]. The notation of *dN* and *dS* was chosen to reflect our focus on the molecular evolution of poliovirus proteins even though we analyze virus populations[77]. To evaluate mutational diversity, two independent measures that quantify diversity were used. First, Shannon’s Entropy, a metric from information theory that quantifies information content, was computed of the mutational space per position as 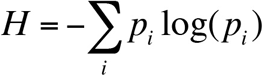, where *p*_*i*_ is the probability of observing mutation *i* amongst the mutations. Second, Simpson’s Index of Diversity 1-D, a biodiversity measure from ecology, was used to assess mutational diversity per position as 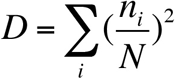, where *n* is the number of mutations of each type *i*, and *N* is the total number of all mutations. For both metrics, diversity profiles of the poliovirus coding sequence, considering all sites with at least 1 mutation, were computed for *Hsp90*_*N*_ and *hsp90*_*i*_. Upon validation that the observed differences between *Hsp90*_*N*_ and *hsp90*_*i*_ are consistent for all passages, average profiles across passages are reported. Global differences in mutation rate, evolutionary rates, and mutational diversity were tested for statistical significance with the Wilcoxon–Mann–Whitney test.

To test how host-cell chaperone levels influence mutations in the virus populations that impact protein folding and stability, we considered three classes of mutations namely destabilizing mutations, mutations that increase the protein aggregation propensity, and mutations that significantly alter sequence hydrophobicity. The effect of all amino acid mutations on protein stability was predicted with FoldX[43] (foldxsuite.crg.es) for poliovirus proteins P1 (PDB structure 2PLV), 3C (4DCD), and 3D (4NLR). The effect of amino acid mutations on protein aggregation propensity was predicted with Tango[44] (tango.crg.es), and changes in amino acid hydrophobicity upon mutation were assessed by the Kyte-Doolittle scale normalized to zero-mean and unitary standard deviation[78]. Mutations were subsequently classified into effect (*E*) and no-effect (*NE*), i.e. destabilizing if the predicted stability effect ΔΔG > 1 kcal/mol, and otherwise non-destabilizing; as aggregation prone if ΔAgg > 2, and otherwise non-aggregation prone; and as hydrophobic if ΔHyd > 4, and otherwise non-hydrophobic else (Suppl. Fig. 4a,b). Solvent accessible surface area (ASA) for each residue was computed from the PDB structures with the DSSP program[79, 80]. Residues were considered buried with ASA < 30, at the interface if the residue contributes ASA > 30 to the monomer interface in the assembled capsid, and as surface residue if ASA > 50 and not at the monomer interface.

To identify sites along the poliovirus sequence that differ significantly in their ability to accommodate destabilizing, aggregation prone, or hydrophobic mutations, we next tested whether the mutation class *E* was statistically associated with the condition of either *Hsp90*_*N*_ or *hsp90*_*i*_ by constructing a 2×2 contingency table for each site composed of the mutation counts of *E_Hsp90*_*N*_, *NE_Hsp90*_*N*_, *E_hsp90*_*i*_, and *NE_hsp90*_*i*_. Finally, sites that are systematically enriched in destabilizing, aggregation prone, or hydrophobic mutations across passages were identified by the Cochran-Mantel-Haenszel (CMH) test on the 2×2×7 contingency table per site and all seven sequenced passages. We found this the most robust approach considering experimental design, the nature of population sequencing and the possibility of spurious adaptation due to laboratory passaging[81]. After correcting p-values with the FDR method[74], significant sites are defined through p < 0.05 and counted *Hsp90*_*N*_ if the odd’s ratio (OR) > 1, and *hsp90*_*i*_ for OR < 1. This identifies sites that are systematically, across passages, under selective pressure to differentially accommodate specific sets of mutations, namely destabilizing, aggregation prone, or hydrophobic mutations as function of host cell chaperone levels.

To identify individual mutations that give rise to significant sites as defined above, Cochran–Mantel–Haenszel test on the 2×2×7 contingency tables were performed for every possible codon mutation along the poliovirus codon sequence comparing the mutation count and reference sequence codon count between *Hsp90*_*N*_ and *hsp90*_*i*_ as well as across passages. After p-value correction with the FDR method, we identified individual significant mutations through p < 0.05, as well as the requirement of them being at a significant site and following the same trend as the significant site, i.e. if a site is enriched in destabilizing mutations at *Hsp90*_*N*_, then the individual mutations at the site also has to be destabilizing and enriched at *Hsp90*_*N*_.

The mutational effect on codon optimality was tested based on the established tRNA adaptation index (tAI)[65] that quantifies how much coding sequences are adapted to the cellular tRNA pool for efficient translation. Here the tAI for HeLa cells based on quantification of tRNA levels through tRNA microarrays[82] was used to compute a host cell tAI. As a control, results were compared to those obtained with a tAI index computed from tRNA gene copy numbers in the human genome[83] (Suppl. Fig. 6). To assess global changes in codon optimality, we computed the ΔtAI = tAI_mutation_ - tAI_reference_ for each sequenced mutation in *Hsp90*_*N*_ and *hsp90*_*i*_ and analyzed the distributions of ΔtAI. To specifically test for an enrichment in mutations to more slowly translated non-optimal codons, we considered the bottom 20% codons with the lowest tAI values as non-optimal. The Cochran–Mantel–Haenszel test was used to identify sequence positions associated with mutations to non-optimal codons at *Hsp90*_*N*_ and *hsp90*_*i*_ across passages.

Data and computer code to reproduce all analyses are available online from https://github.com/pechmann/polio.

## Acknowledgments

We thank members of the Frydman and Andino labs for discussions and Drs. Adam Lauring, Marco Vignuzzi, Charles Parnot and Marcus Feldman for advice and comments. This work was financially supported by NIH grants to RA and JF and a NIH/NIAID Postdoctoral Fellowship to AA. SP holds the Canada Research Chair in Computational Systems Biology. RG holds the Ramon y Cajal fellowship from the Spanish Ministry of Economy and Competitiveness (RYC-2015-17517).

## Author Contributions

RG, JF and RA designed the study; RG, SP, JF and RA interpreted the data and wrote the manuscript. RG carried out all experiments and SP the computational analyses; AA performed the CirSeq sequencing; RA and JF contributed equally to the study.

## Supplementary Figures

**Supplementary Fig. 1:**
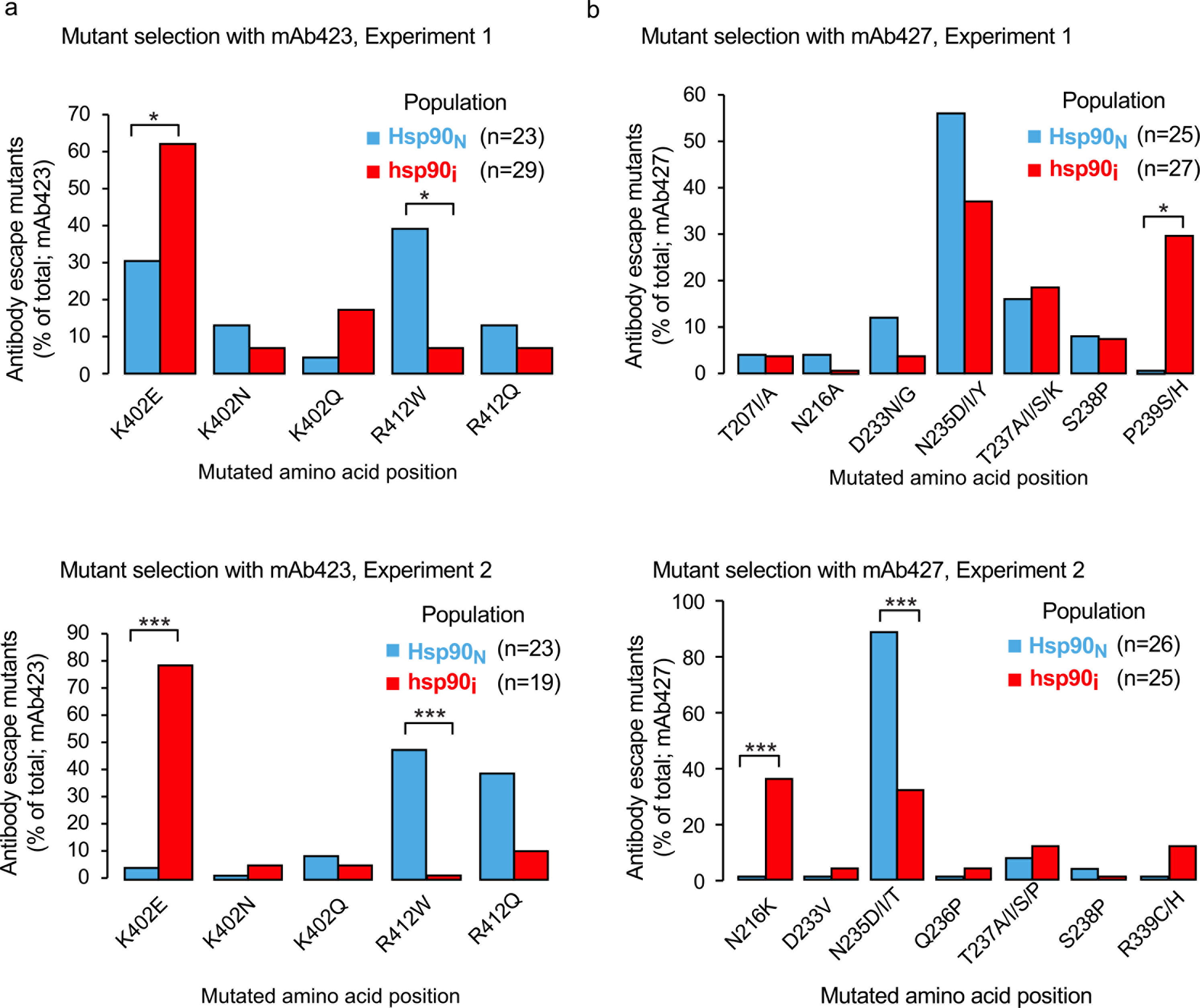
Results of individual mutant capsid selection assays. (**a**,**b**) Abundance and distribution of mAb423 (a) or mAb427 (b) escape variants in populations of Sabin 1 (a) or WT (b) poliovirus from *Hsp90*_*N*_ and *hsp90*_*i*_ populations from 2 separate experiments. The number of mAb-resistant virus variants examined (n) is indicated. * p < 0.05, ** p < 0.01, *** p < 0.005 by 2 tail Fisher’s exact test.

**Supplementary Fig. 2:**
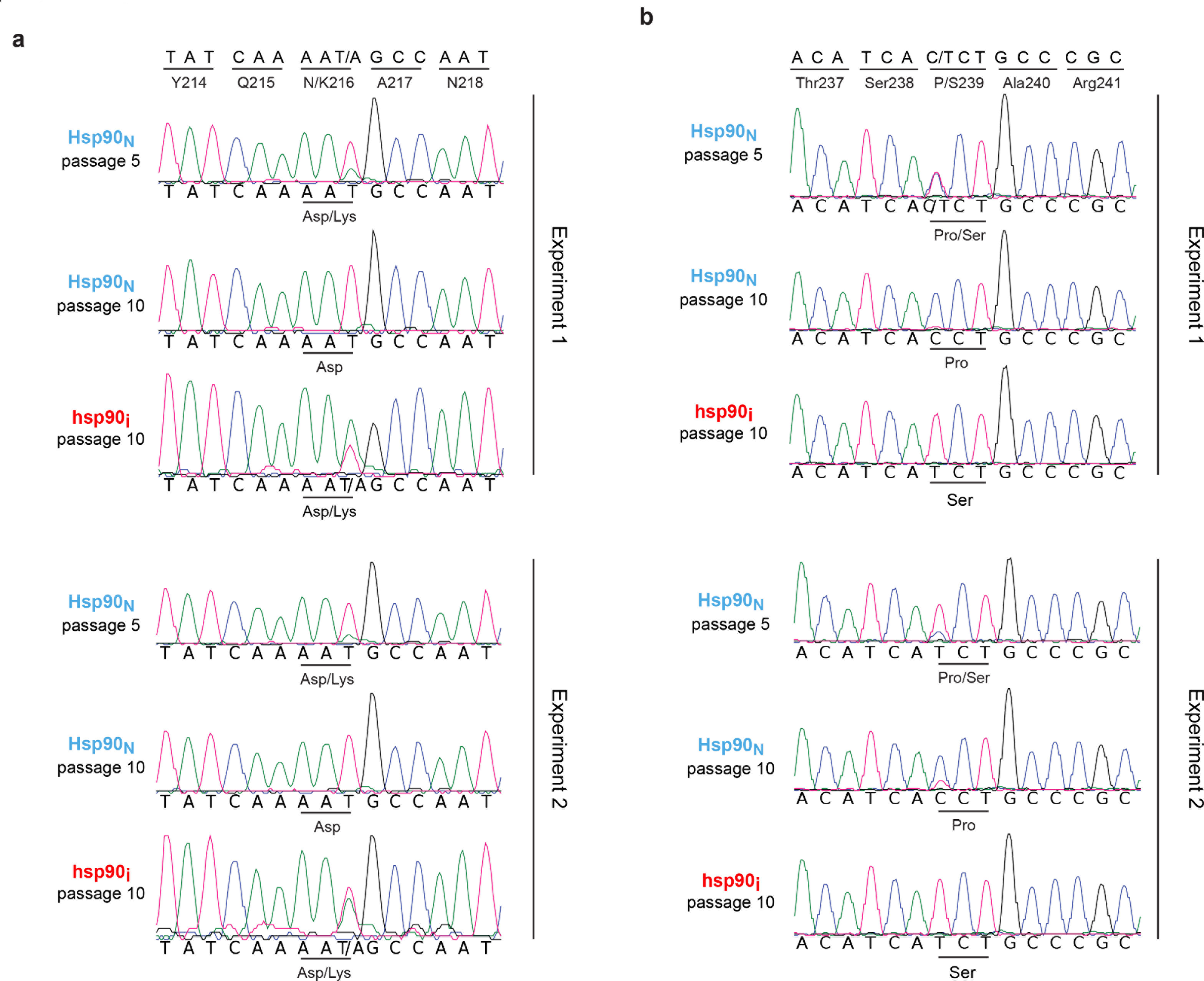
Sanger sequencing analysis of competition experiments showing effect of Hsp90 activity on viral fitness of escape variants. (**a**,**b**) Results of competition experiments testing the fitness of N216K (a) or P239S (b) variants under conditions of normal or low Hsp90 activity. An initial infection was performed with a 1:5 ratio to wildtype (N216 or P239) to *hsp90*_*i*_ variant (K216 or S239) virus. Viruses were then passaged for a total of 10 passages and competition results assessed by Sanger sequencing the passage 10 (p10) virus populations. Intermediate passage 5 (p5) results for normal Hsp90 conditions showing mixed population are also shown.

**Supplementary Fig. 3:**
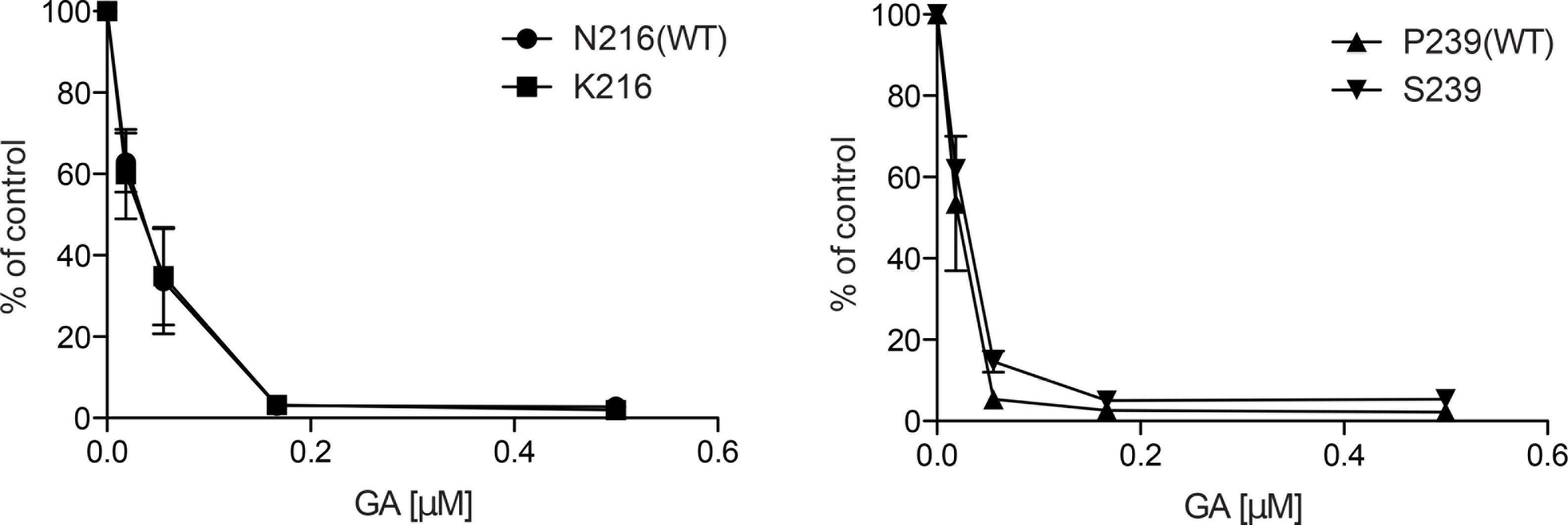
mAb escape variants from *hsp90*_*i*_ populations do not show reduced sensitivity to Hsp90 inhibition. (**a**) Percent of virus production relative to no Hsp90 inhibitor for both WT and *hsp90*_*i*_ variants K216 and S239 observed to be enriched in mAb selection experiments. Results represent data from 3 experiments, with mean and SEM plotted. No significant differences were observed (p > 0.1 by MWW test).

**Supplementary Fig. 4:**
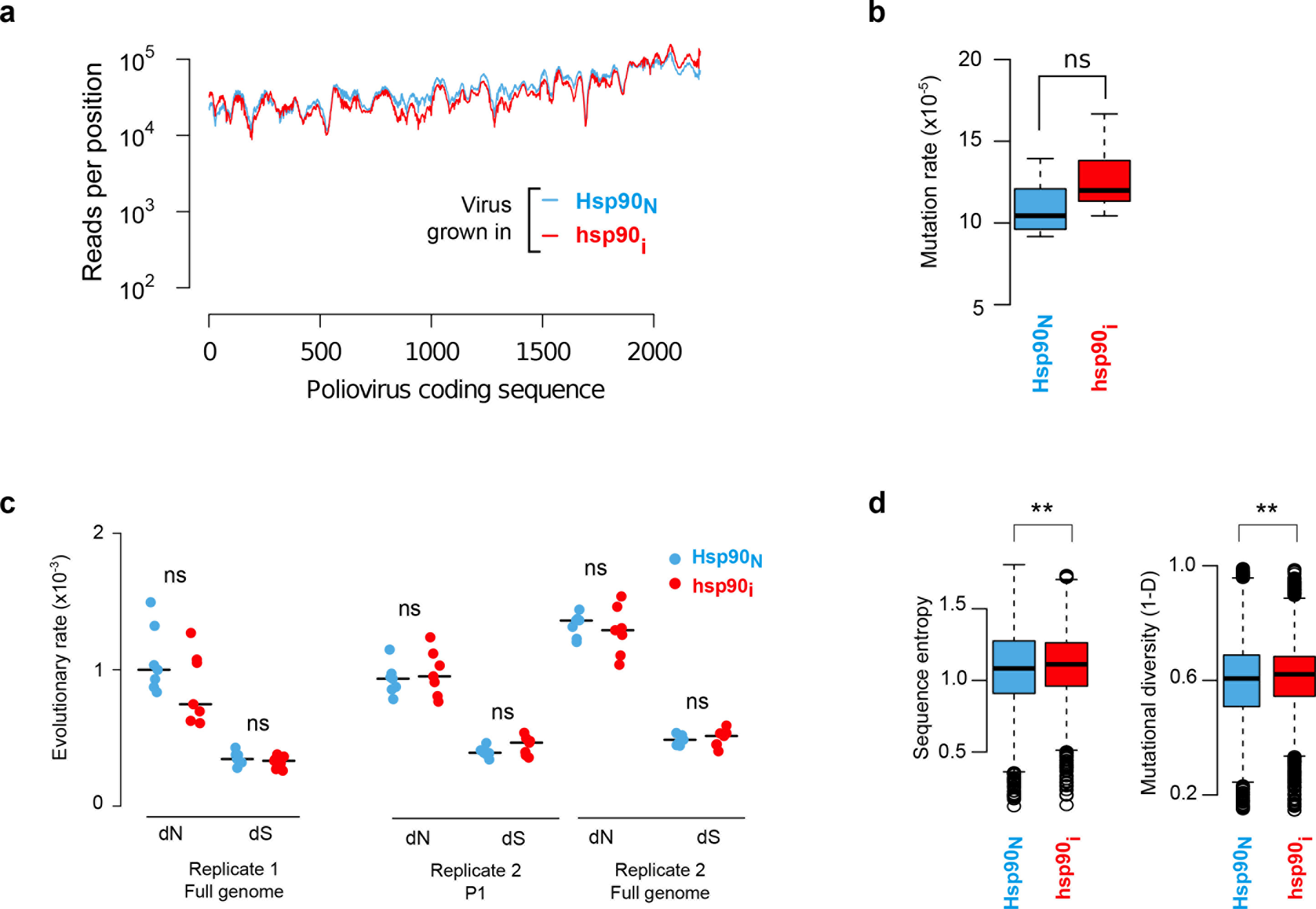
Reproducibility of CirSeq analysis. (**a**) CirSeq sequencing coverage of the poliovirus coding region for Replica 2. (**b**) Mutation rate of virus populations in Replica 2. No significance difference is observed for viral passages between *Hsp90*_*N*_ and *hsp90*_*i*_ conditions (p = 0.165 by MWW test). (**c**) Evolutionary rates *dN* and *dS* of poliovirus populations. No significant differences in the standard evolutionary rates of nonsynonymous (*dN*) and synonymous (*dS*) substitutions between virus populations from *Hsp90*_*N*_ and *hsp90*_*i*_ conditions were observed for the full coding sequence of Replica 1 (*dN*: p = 0.38, *dS*: p = 0.26 by MWW test), the capsid protein P1 of Replica 2 (*dN*: p = 0.26, *dS*: p = 0.71 by MWW test), or the full coding sequence of Replica 2 (*dN*: p = 0.71, *dS*: p = 0.62 by MWW test).(**d**) Mutational diversity of poliovirus populations for Replica 2. Viral populations passaged under reduced Hsp90 activity (*hsp90*_*i*_) show higher diversity of mutations as estimated by sequence entropy (p = 0.0098 by MWW test) or mutational diversity (1-*D*), with *D* as Simpson’s diversity index (p = 0.0011 by MWW test). Significance levels are indicated by ns for not significant and ** for p < 0.01.

**Supplementary Fig. 5:**
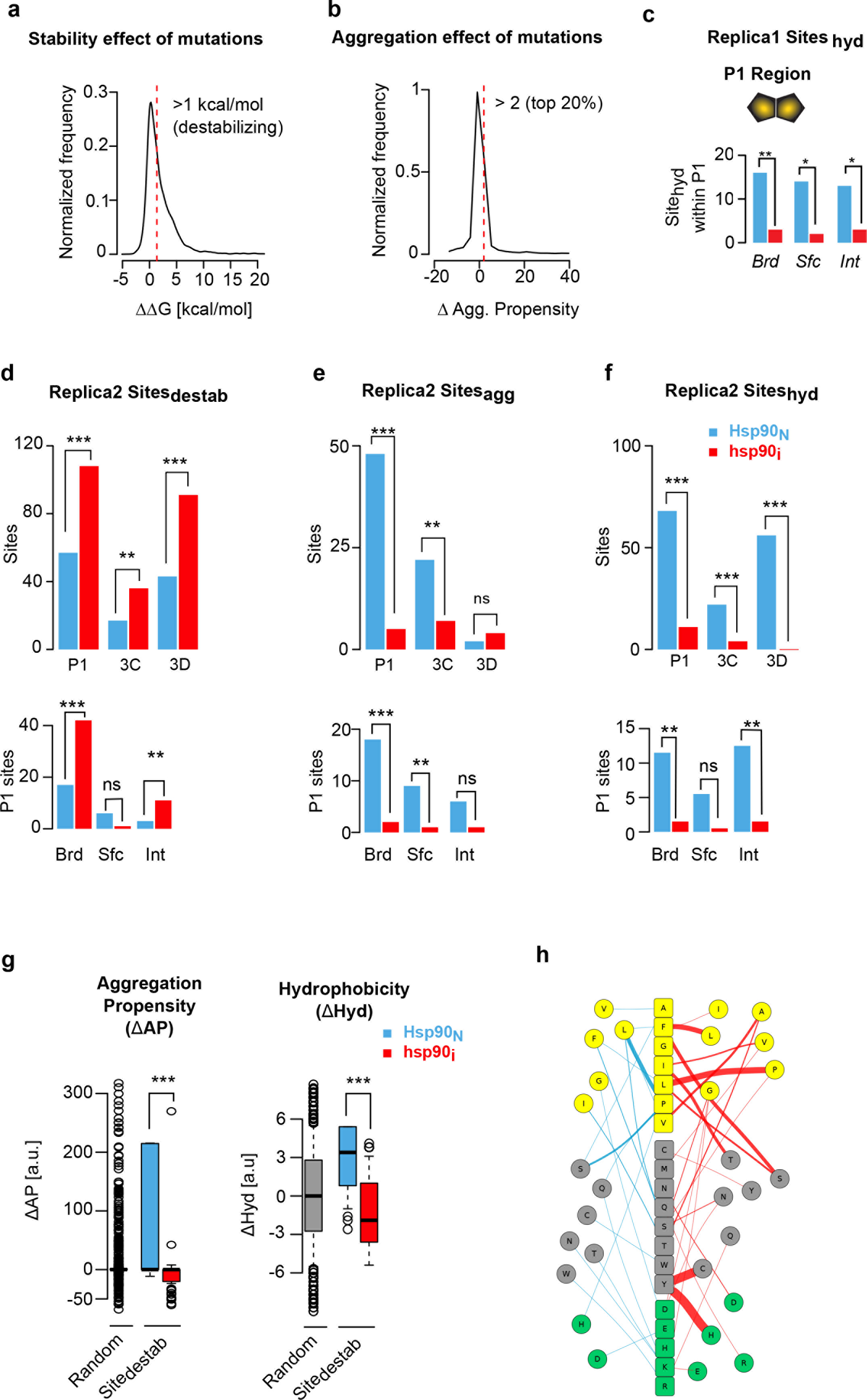
Distribution of stability and aggregation effects of variants and reproducibility of variants analyses for replica 2. (**a**) Distribution of the stability effect of all possible amino acid mutations in poliovirus proteins P1, 3C protease and 3D polymerase. Mutations were calculated using the FoldX algorithm and those having a ΔΔG > 1kcal/mol are considered destabilizing. (**b**) Distribution of the effect of all possible amino acid mutations in poliovirus on the aggregation propensity predicted with the TANGO algorithm. We classified the top 20%,i. e. mutations that increase the aggregation propensity by more than 2 [a.u.] as aggregation prone mutations. (**c**) The distribution of Site_hyd_ at buried (Brd), interface (Int), or surface (Sfc) positions across P1 based on the capsid crystal structure (PDB: 2PLV) for Replica 1. (**d**) Sites that differ in their ability to accommodate destabilizing variants between *Hsp90*_*N*_ and *hsp90*_*i*_ conditions (Site_destab_) for Replica 2. All analyzed viral proteins accommodate more Sitesdestab at lower Hsp90 activity (P1: p<10^-4^, 3C: p = 0.0061, 3D: p <10^-5^ by MWW test). Within P1, Sitesdestab differ significantly in buried (Brd; p = 0.00078 by Fisher’s test) and interface (Int; p = 0.0052 by Fisher’s test) position, whereas no significant differences were observed for the capsid surface (Srf; p = 0.12 by Fisher’s test). (**e**) Sites that differ in their ability to accommodate aggregation prone mutations between *Hsp90*_*N*_ and *hsp90*_*i*_ conditions (Sites_agg_) for Replica 2. P1 and 3C accommodate significantly fewer Sites_destab_ at lower Hsp90 activity (P1: p<10^-9^, 3C: p = 0.0042, 3D: p = 0.62 by MWW test). Within P1, Sites_agg_ differ significantly at buried and surface positions (buried: p= 0.00014, interface: p = 0.06, surface: p = 0.007 by Fisher’s test). (**f**) Sites that differ in their ability to accommodate hydrophobic mutations between *Hsp90*_*N*_ and *hsp90*_*i*_ conditions (Sites_hyd_) for Replica 2. All analyzed viral proteins accommodate significantly fewer Sites_hyd_ at lower Hsp90 activity (P1: p < 10^-11^, 3C: p = 0.0017, 3D: p = < 10^-6^ by MWW test). Within P1, Sites_hyd_ differ significantly at buried and interface positions across the P1 protein (buried: p = 0.0058, interface: p = 0.0029, surface: p = 0.06 by Fisher’s test). (**g**) Mutational pleiotropy in Replica 2. Destabilizing mutations enriched in *Hsp90*_*N*_ are significantly more aggregation prone (p = 0.0006 by MWW test) and hydrophobic (p = 6.85 × 10^-8^ by MWW test) compared to destabilizing mutations enriched in *hsp90*_*i*_. (**h**) Destabilizing variants observed in Site_destab_ in *Hsp90*_*N*_ and *hsp90*_*i*_ populations of replica 2. The edge weight scales with the number of occurrences of variants enriched in *Hsp90*_*N*_ (left) and *hsp90*_*i*_ (right), respectively, and the WT amino acid is indicated in the middle. Significance levels are indicated by ns for not significant, ** for p < 0.01, and *** for p < 0.001.

**Supplementary Fig. 6:**
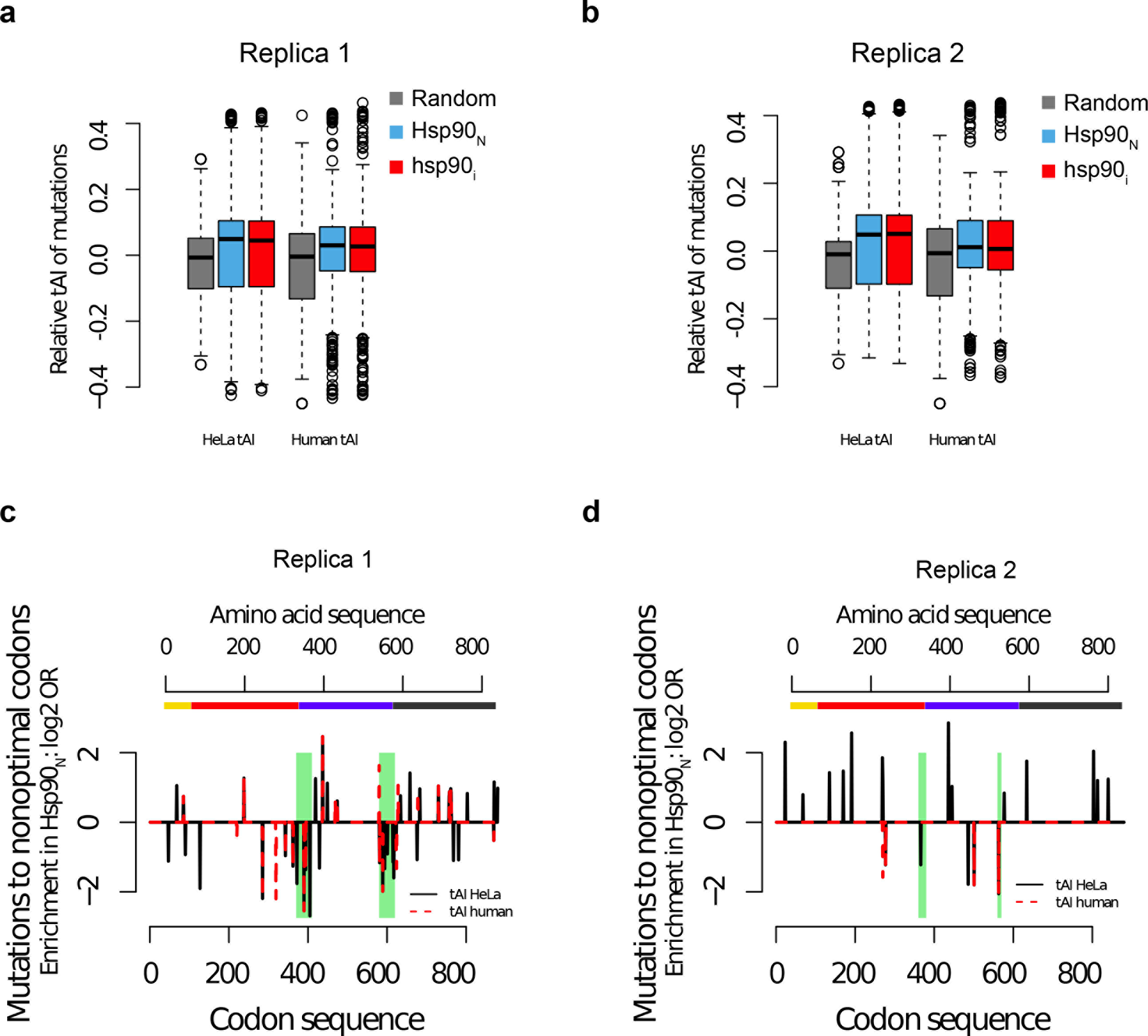
Analysis of codon optimality for *Hsp90*_*N*_ and *hsp90*_*i*_ populations. (**a**,**b**) Relative tRNA adaptation index (tAI) of sequence mutations based on either human or HeLa cell tAI for both replica 1 (a) or 2 (b). (**c**,**d**) Sites showing significant codon deoptimization differences between *Hsp90*_*N*_ and *hsp90*_*i*_ populations in replica 1 (c) or replica (2).

